# Which are major players, canonical or non-canonical strigolactones?

**DOI:** 10.1101/195255

**Authors:** Koichi Yoneyama, Xiaonan Xie, Kaori Yoneyama, Takaya Kisugi, Takahito Nomura, Yoshifumi Nakatani, Kohki Akiyama, Christopher S. P. McErlean

## Abstract

Strigolactones (SLs) can be classified into two structurally distinct groups: canonical and non-canonical SLs. Canonical SLs contain the ABCD ring system, and non-canonical SLs lack the A, B, or C ring but have the enol ether–D ring moiety which is essential for biological activities. The simplest non-canonical SL is the SL biosynthetic intermediate carlactone (CL). In plants, CL and its oxidized metabolites such as carlactonoic acid and methyl carlactonoate, are present in root and shoot tissues. In some plant species including black oat (*Avena strigosa*), sunflower (*Helianthus annuus*), and maize (*Zea mays*), non-canonical SLs are major germination stimulants in the root exudates. Various plant species such as tomato (*Solanum lycopersicum*) release carlactonoic acid, and poplar (*Populus* spp.) was found to exude methyl carlactonoate into the rhizosphere. These results suggest that both canonical and non-canonical SLs are active as host recognition signals in the rhizosphere. In contrast, limited distribution of canonical SLs in the plant kingdom and structure- and stereo-specific transportation of canonical SLs from roots to shoots suggest that plant hormones inhibiting shoot branching are not canonical SLs but are rather non-canonical SLs.

**Abbreviations**
AM: arbuscular mycorrhizal
CL: carlactone
CLA: carlactonoic acid
4DO: 4-deoxyorobanchol
5DS: 5-deoxystrigol
GC–MS: gas-chromatography-mass spectrometry
LC–MS/MS: liquid chromatography-tandem mass spectrometry
MeCLA: methyl carlactonoate
NMR: nuclear magnetic resonance
SL: strigolactone

Supplemental files: Synthesis of 7-hydroxy-5-deoxystrigol stereoisomers, spectroscopic data of synthetic compounds and the natural stimulant in dokudami root exudates.

**Scheme S1.** Synthesis of racemic mixture of 7α- and 7β-hydroxy-5-deoxystrigol.

**Fig. S1.** ^1^H NMR spectrum of natural stimulant in dokudami root exudates.

**Fig. S2.** LC–MS/MS chromatograms of synthetic standards and natural stimulant.

**Fig. S3.** LC–MS/MS chromatograms of synthetic 7β-hydroxy-5-deoxystrigol and natural stimulant.

**Fig. S4.** LC–MS/MS chromatograms of synthetic 7β-hydroxy-5-deoxystrigol and natural stimulant.

**Highlight:** The chemistry of canonical and non-canonical strigolactones and their distribution in the plant kingdom are summarized in relation to their biological activities in the rhizosphere and in plants.

## Introduction

Natural strigolactones (SLs) are carotenoid-derived plant hormones or their precursors that regulate plant growth and developmental processes through cross-talk with other hormones and also participate in plant–root parasitic plants and plant–microbe interactions in the rhizosphere (Xie *et al.*, 2010; Al-Babili and Bouwmeester, 2015). SLs can be classified into two groups based on their chemical structures. Strigol (**1**) and related compounds that contain the ABC ring system connected to the methylbutenolide D ring via an enol ether bridge are called canonical SLs (Al-Babili and Bouwmeester, 2015). So far, 23 canonical SLs have been characterized from plant root exudates through bioassay-guided purifications (Xie *et al.*, 2010; Xie, 2016). Most were isolated as germination stimulants for root parasitic weeds, except for 5-deoxystrigol (**3**) which was originally identified as the hyphal branching factor for arbuscular mycorrhizal (AM) fungi (Akiyama *et al.*, 2005) (**Fig. 1**).

**Fig. 1.**
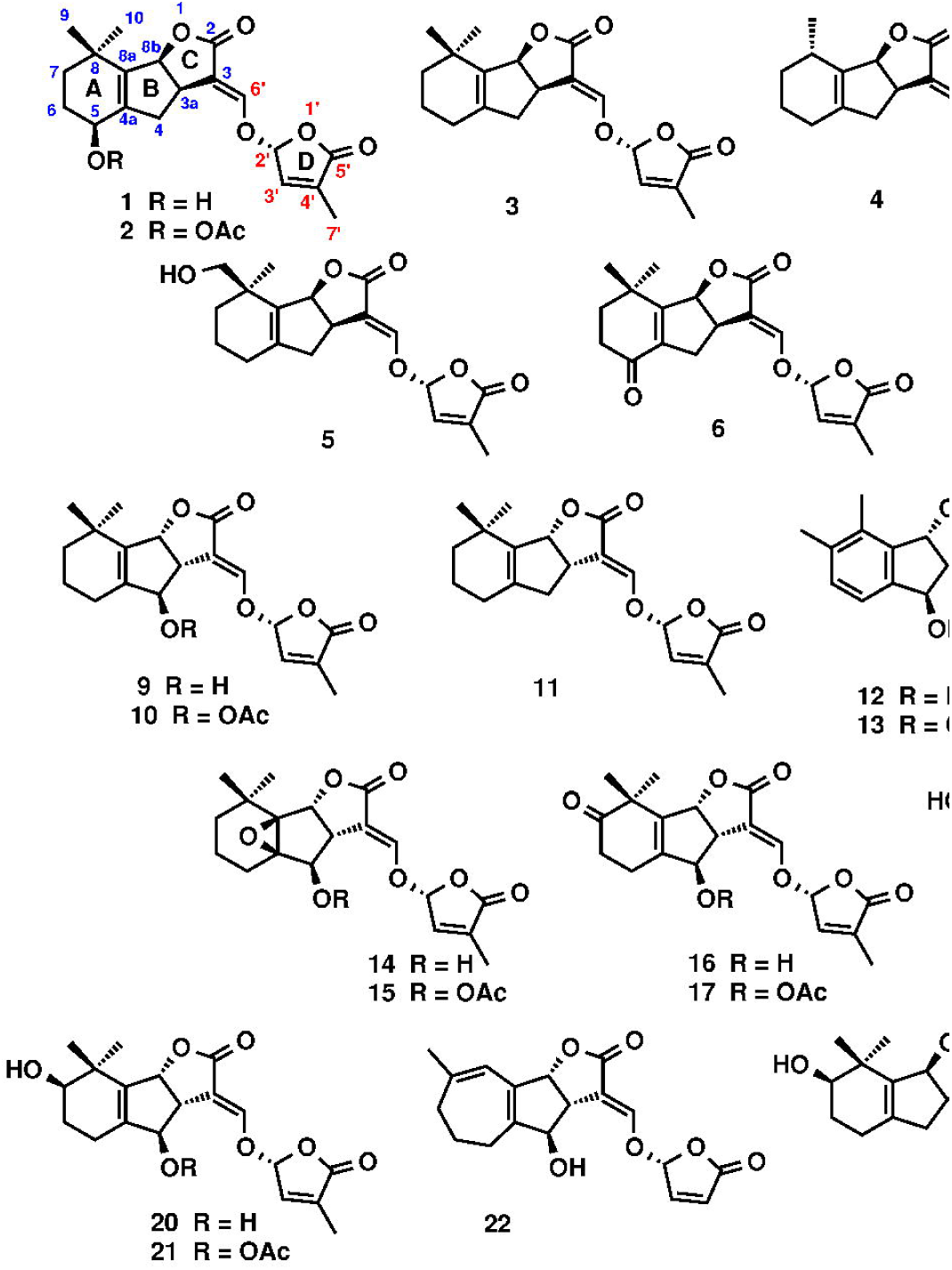
Structures of canonical strigolactones. Strigol-type (**1**–**8**, **23**) and orobanchol-type strigolactones (**9**–**22**).

Extensive studies on the structure–activity relationships of SLs showed that the enol ether–D ring moiety was essential for biological activities (Yoneyama *et al.*, 2009; Zwanenburg *et al.*, 2009; Akiyama *et al.*, 2010; Yoneyama *et al.*, 2010; Boyer *et al.*, 2012). In addition, stereochemistry, in particular at the 2’-position of the D ring, is important for high potencies (Thuring *et al.*, 1997; Flematti *et al.*, 2016). The introduction of a hydroxyl group on the A or B ring generally reduces stability and thus germination stimulation of *Striga* spp. (Gobena *et al.*, 2017) but not *Orobanche* spp. (Kim *et al.*, 2010). Various synthetic SL mimics carrying the essential structural unit have been developed (Nefkens *et al.*, 1997; Kondo *et al.*, 2007; Zwanenburg *et al.*, 2009; Cohen *et al.*, 2013; Fukui *et al.*, 2013; Boyer *et al.*, 2014; Screpanti *et al.*, 2016).

Recently, germination stimulants lacking the ABC ring structure but containing the D ring were characterized and are called non-canonical SLs. They are zealactone (methyl zealactonoate, **24**) (Charnikhova *et al.*, 2017; Xie *et al.*, 2017), avenaol (**25**) (Kim *et al.*, 2014), heliolactone (**26**) (Ueno *et al.*, 2014), and two novel stimulants (**27** and **28**). The simplest non-canonical SL is carlactone (CL, **29**) (Alder *et al.*, 2012), a SL biosynthetic precursor. Its oxidized metabolite, carlactonoic acid (CLA, **30**) (Abe *et al.*, 2014), has been detected in root exudates from various plant species (Yoneyama *et al.*, 2017). Some species such as poplar (*Populus* spp.) release methyl carlactonoate (MeCLA, **31**) into the rhizosphere, which can interact with the SL receptor D14 (**Fig. 2**) (Abe *et al.*, 2014; Yoneyama *et al.*, 2017). These results indicate that, in addition to canonical SLs, non-canonical SLs including CLA and MeCLA are likely involved in chemical communications in the rhizosphere (Yoneyama *et al.*, 2017).

**Fig. 2.**
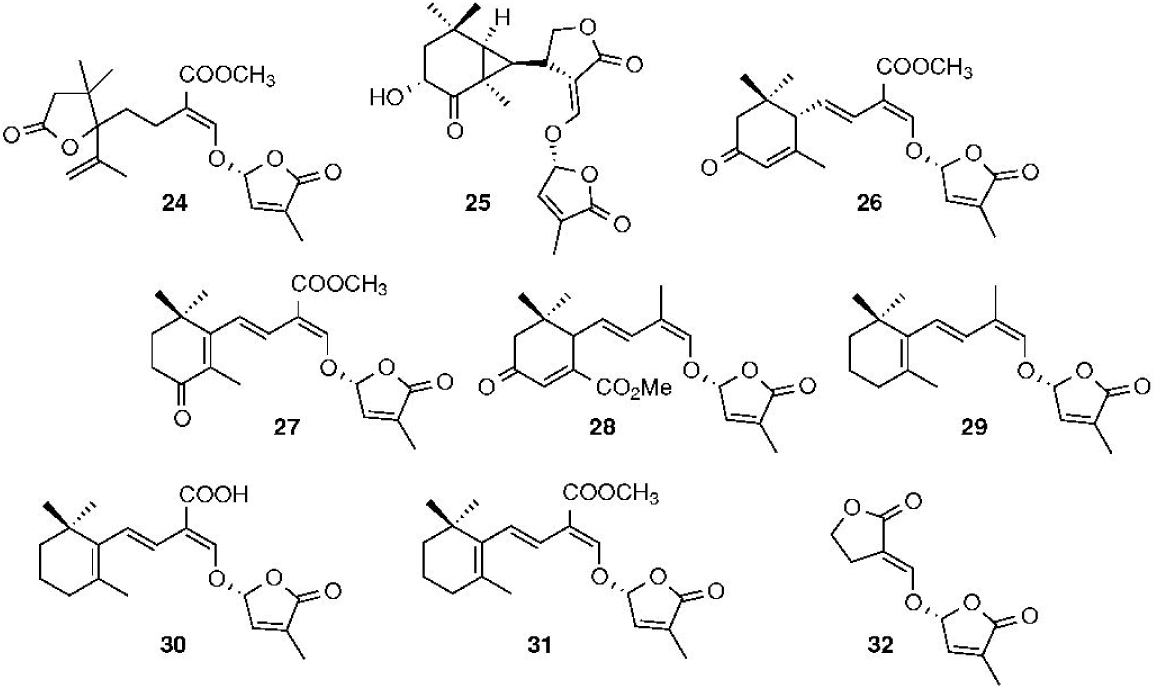
Structures of non-canonical strigolactones and GR5 (**32**)

In this review, chemistry of canonical and non-canonical SLs and their distribution in the plant kingdom are summarized and discussed in relation to their involvement in parasite seed germination, AM fungi colonization, and shoot branching inhibition.

## Canonical SLs

The canonical SLs characterized to date are strigol (**1**), strigyl acetate (**2**), 5-deoxystrigol (5DS, **3**), sorgolactone (**4**), sorgomol (**5**), strigone (**6**), 4-hydroxy-5-deoxystrigol (*ent*-2’ -*epi*-orobanchol, **7**), 4-acetoxy-5-deoxystrigol (*ent*-2’-*epi*-orobanchyl acetate, **8**), orobanchol (**9**), orobanchyl acetate (**10**), 4-deoxyorobanchol (4DO, **11**), solanacol (**12**), solanacyl acetate (**13**), fabacol (**14**), fabacyl acetate (**15**), 7-oxoorobanchol (**16**), 7-oxoorobanchyl acetate (**17**), 7α-hydroxyorobanchol (**18**), 7α-hydroxyorobanchyl acetate (**19**), 7β-hydroxyorobanchol (**20**), 7β-hydroxyorobanchyl acetate (**21**), and medicaol (**22**) (Cook *et al.*, 1969; Hauck *et al.*, 1992; Yokota *et al.*, 1998; Akiyama *et al.*, 2005; Xie *et al.*, 2007; Umehara *et al.*, 2008; Xie *et al.*, 2008; Xie *et al.*, 2009a; Xie *et al.*, 2009b; Chen *et al.*, 2010; Kisugi *et al.*, 2013; Tokunaga *et al.*, 2015; Xie, 2016). According to C ring stereochemistry, these canonical SLs can be divided into two groups: strigol- (**1**–**8**) and orobanchol-type SLs (**9**–**22**), which have a β- and an α-oriented C ring, respectively (**Fig. 1**) (Xie *et al.*, 2013; Al-Babili and Bouwmeester, 2015). In general, plants appear to produce either strigol- or orobanchol-type SLs as major SLs; however, some species such as tobacco (*Nicotiana tabacum*) produce both (Xie *et al.*, 2013). The three gymnosperms so far examined – Japanese black pine (*Pinus thunbergii*), cedar (*Cryptomeria japonica*), and gingko (*Gingko biloba*) – and the lycophyte spike moss (*Selaginella moellendorfii*) produce orobanchol-type SLs (Yoneyama *et al.*, 2017) (**Table 1**). Many angiosperms are orobanchol-type SL producers but cotton (*Gossypium hirsutum*) (Cook *et al.*, 1969; Sato *et al.*, 2005) and strawberry (*Fragaria vesca*) plants exude only strigol-type SLs (Xie *et al.*, unpublished). Chinese milk vetch (*Astragalus membranaceus*) is unique among the Fabaceae plants as it produces strigol-type SLs (Yoneyama *et al.*, 2008). Previously, we reported that Chinese milk vetch produced orobanchyl acetate but re-examination with our new liquid chromatography–tandem mass spectrometry (LC–MS/MS) system suggests that the signal assigned to orobanchyl acetate may be background noise. We encountered similar problems in SL detection from *Physcomitrella patens* (Proust *et al.*, 2011). Although we detected CL in *P. patens* gametophores as expected, we could not detect any known SLs with the new LC–MS/MS system. The Poaceae family contains both a strigol-type SL producer, sorghum (*Sorghum bicolor*) (Yoneyama *et al.*, 2007; Xie *et al.*, 2008), and an orobanchol-type SL producer, rice (*Oryza sativa*) (Umehara *et al.*, 2008). Recently, some *Striga*-resistant sorghum cultivars were shown to produce orobanchol, which is a weak *Striga* germination stimulant but a potent branching factor for AM fungi. These cultivars are thus resistant to *Striga* but still good hosts for AM fungi (Gobena *et al.*, 2017). We also observed that despite the distinctive difference in SL composition, the levels of AM colonization and the community compositions did not differ between the *Striga*-susceptible and -resistant maize cultivars (Yoneyama *et al.*, 2015). Identities of SLs in these maize cultivars need to be confirmed as these data were obtained with our old MS system.

**Table 1.**
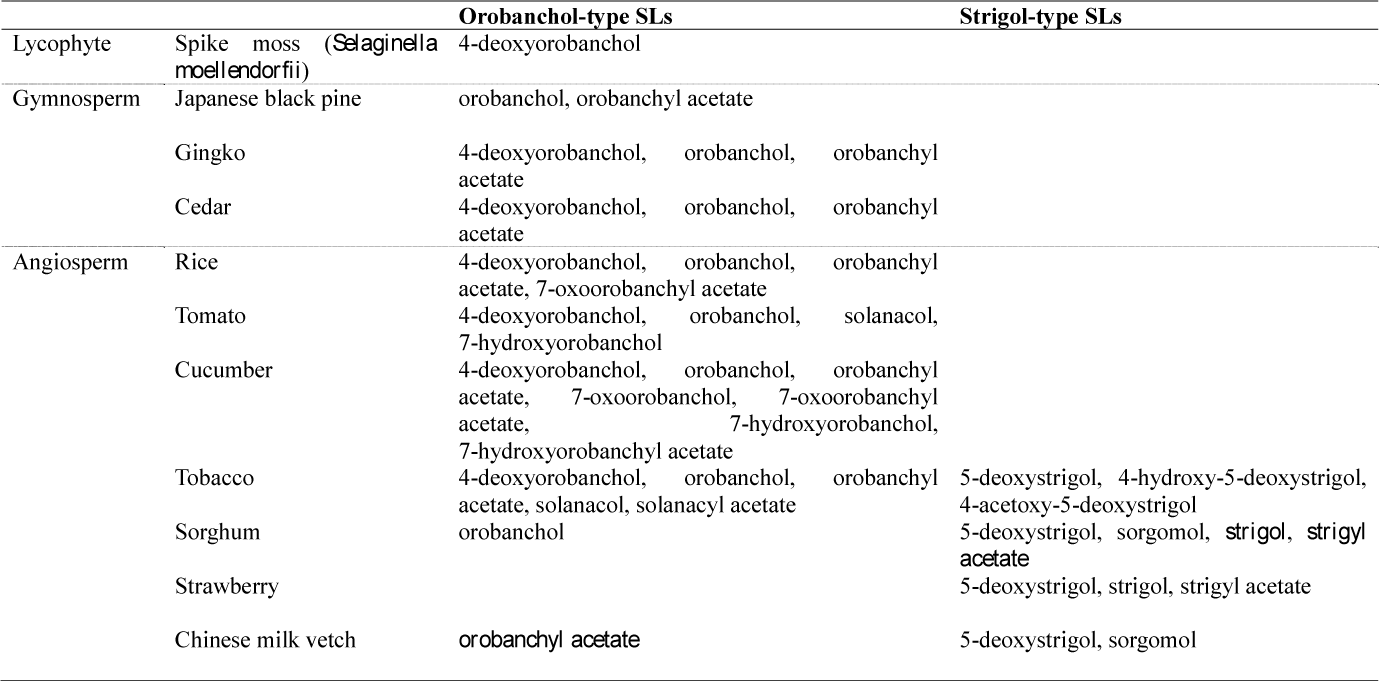
Distribution of canonical strigolactones (SLs) in the plant kingdom. SLs in italics need to be confirmed.

In addition to these 22 canonical SLs, we characterized at least three novel canonical SLs, 7β-hydroxy-5-deoxystrigol (**23**) from dokudami (*Houttuynia cordata*) and two from tall goldenrod (*Solidago altissima*), indicating that the number of canonical SLs in the plant kingdom may increase as we examine more species. The structures of the two canonical SLs produced by tall goldenrod will be reported elsewhere.

It is intriguing to understand why plants produce orobanchol- and strigol-type SLs. One of rice MORE AXILLARY GROWTH 1 (MAX1) orthologs, Os900 expressed in yeast converts CL to 4DO but not to 5DS (Zhang *et al.*, 2014), presumably via CLA and 18-hydroxy-CLA *in vitro* (Yoneyama *et al.*, 2017). It is likely that Low Germination Stimulant 1 (LGS1), a sulfotransferase, is involved in formation of strigol-type SLs (Gobena *et al.*, 2017) and therefore orobanchol-type SLs may have evolved due to mutations in the *LGS1* gene, as observed for *Striga*-resistant sorghum cultivars. However, the wide distribution of the orobanchol-type SL, 4DO, in gymnosperms, angiosperms, and a lycophyte may imply that orobanchol-type SLs evolved earlier. The distribution of canonical SLs, orobanchol- and strigol-type SLs, in the root exudates from various plant species is listed in **Table 1**.

### 7β-Hydroxy-5-deoxystrigol from dokudami root exudate

Dokudami is a Chinese medicinal plant commonly found in Japan. Strigol (**1**) and strigone (**6**) have been identified as a major and a minor SL, respectively, in its root exudate which contained several SL-like compounds as minor germination stimulants (Kisugi *et al.*, 2013). Three had the molecular formula C_19_H_22_O_6_ but their retention times in LC–MS/MS analyses differed from those of known SLs, such as strigol. They had quite low contents in root exudate, ca. 1/500 that of strigol, and we isolated a very small amount of one (< 2 µg) with some impurities. The MS spectroscopic data and the proton nuclear magnetic resonance (NMR) spectrum of this stimulant suggested that it was a mono-hydroxy-canonical SL, most likely carrying a hydroxyl group at C_7_. Comparing retention times of the natural stimulant and synthetic standards on C_18_, NAP, and chiral columns using LC–MS/MS, showed that the structure was 7β-hydroxy-5-deoxystrigol (**23**). Synthesis and spectroscopic data of both natural and synthetic 7β-hydroxy-5-deoxystrigol and its isomers are included in supplementary data.

## Non-canonical SLs

Although maize (*Zea mays*), a major host of *Striga*, had been reported to exude strigol (Siame *et al.*, 1993), (Jamil *et al.*, 2012) detected neither strigol nor any other canonical SL in maize root exudates with LC–MS/MS analysis. Instead, they found two novel germination stimulants, SL1 and SL2. Similarly, root exudates from black oat (*Avena strigosa*), which strongly induced *Striga* and *Orobanche* germination, did not contain known canonical SLs. Therefore, germination stimulants produced by maize and black oat have been examined extensively, and two non-canonical SLs, zealactone (**24**) (Charnikhova *et al.*, 2017; Xie *et al.*, 2017) and avenaol (**25**) (Kim *et al.*, 2014) were isolated from their root exudates, respectively, and their structures determined. Very recently, the stereochemical structure of avenaol was confirmed by total synthesis (Yasui *et al.*, 2017). These non-canonical SLs lack the A, B, or C ring but retain the enol ether–D ring moiety which is essential for biological activities of SLs. Maize was found to produce several non-canonical SLs, and SL2 was determined to be zealactone (Charnikhova *et al.*, 2017; Xie *et al.*, 2017). SL1 was found to be mixtures of at least two isomers which were difficult to separate. Another non-canonical SL, heliolactone (**26**), was isolated from root exudates of sunflower (*Helianthus annuus*), the host of *O. cumana*. Sunflower plants do not produce known canonical SLs (Ueno *et al.*, 2014). In addition to these non-canonical SLs, we isolated at least two non-canonical SLs, one (**27**) from rice and the other (**28**) from black oat root exudates (**Fig. 2**). Compound **27** is one of the putative methoxy-5DS isomers produced by rice plants (Jamil *et al.*, 2011). The structures of **27** and **28** in **Fig. 2** are only tentative and need to be confirmed, preferably by comparing spectroscopic data with those of synthetic standards. This is because these compounds are present in root exudates in quite low amounts and, in particular, **27** occurs as a mixture of isomers which hampers further purification. Furthermore, these non-canonical SLs are less stable than canonical SLs and gradually decompose during purification and storage.

The first reported non-canonical SL was CL (**29**), a SL biosynthetic precursor (Alder *et al.*, 2012). The levels of CL in plant tissues appear to be rather high compared to canonical SLs and are not affected by phosphate starvation, indicating that response to phosphate starvation is regulated in the SL biosynthetic steps after CL formation (Seto *et al.*, 2014). Although CL is rather lipophilic, it was detected in the root exudate of tall goldenrod (Xie *et al.*, unpublished). The oxidized metabolite of CL, CLA (**30**) (Abe *et al.*, 2014), has been detected in root exudates from various plant species. Some plant species such as poplar (*Populus* spp.) release MeCLA (**31**) (Yoneyama *et al.*, 2017). These results indicate that CLA and MeCLA are likely involved in chemical communications in the rhizosphere. The model legume plant *Lotus japonicus* exudes the simplest canonical SL, 5DS (Akiyama *et al.*, 2005), and also a non-canonical SL, tentatively named lotuslactone (methyl lotuslactonoate), whose structure will be reported elsewhere.

So far, less than ten non-canonical SLs have been characterized. It is likely, however, that many more will be identified, since they can be structurally more diverse than canonical SLs. In addition, we have only recently understood how to handle and protect non-canonical SLs from degradation during isolation and purification.

Most canonical SLs are C_19_ compounds except for acetoxyl derivatives such as orobanchyl acetate which contain additional two carbons, and sorgolactone, a C_18_ SL. In contrast, non-canonical SLs contain an additional carbon atom, except for CL and CLA, and thus are C_20_ compounds. It is likely that this additional carbon comes from the ester methyl group and therefore these non-canonical SLs appear to be derived from MeCLA or its isomers and their hydroxyl derivatives. Possible biosynthetic pathways for non-canonical SLs are shown in **Fig. 3**. Non-canonical SLs containing a methoxycarbonyl group should be named methyl esters rather than lactones as in the case of MeCLA. In addition, their corresponding free acids may occur in plants and in their root exudates. However, for example, zealactone is much easier to pronounce and remember than methyl zealactonoate. The corresponding free acids for zealactone and heliolactone should be named zealactonoic acid and heliolactonoic acid, respectively.

**Fig. 3.**
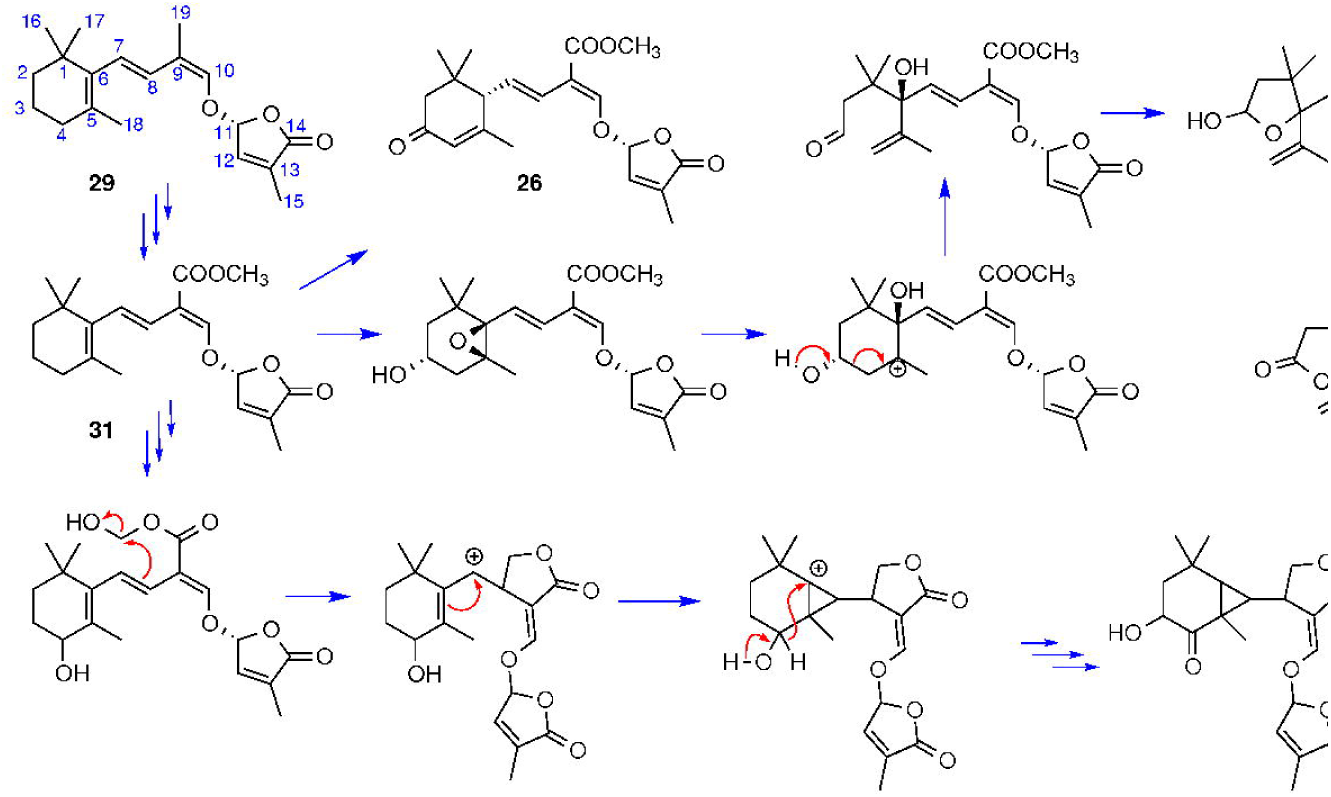
Possible synthetic pathways for non-canonical strigolactones.

Although both canonical and non-canonical SLs are chemically unstable, they may persist in the slightly acidic rhizosphere longer than would be expected in bulk soil (Bertin *et al.*, 2003). In general, canonical SLs appear to be slightly more stable than non-canonical SLs. For example, except for 7-hydroxyorobanchols (**18**, **20**), canonical SLs can be stored safely after organic solvents are completely evaporated. In contrast, it is better to store non-canonical SLs as solutions in organic solvents because even CL, the most stable non-canonical SL, decomposes very rapidly when concentrated. However, we detected highly unstable MeCLA in some samples sent from overseas, indicating that non-canonical SLs may be more stable when included in a mixture of various plant primary and secondary metabolites (Bertin *et al.*, 2003). All of the plant species listed in **Table 1** have been confirmed to exude CLA (Yoneyama *et al.*, 2017).

In addition to the canonical and non-canonical SLs shown in **Figs 1** and **2**, Solanaceae plants, tomato, eggplant (*Solanum melongena*), potato (*Solanum tuberosum*), sweet pepper (*Capsicum annuum*), habanero (*Capsicum chinense*), and tobacco, exude four, four, five, five, and six isomers of putative didehydro-orobanchol, respectively, whose structures remain to be characterized.

## SLs as plant hormones regulating shoot branching

In 2008, two research groups independently identified SLs or their further metabolites as a novel class of plant hormones inhibiting shoot branching or axillary bud outgrowth (Gomez-Roldan *et al.*, 2008; Umehara *et al.*, 2008). Since then, despite extensive studies, these true plant hormones have not unequivocally been identified.

This hormone was first suggested by Christine Beveridge through the forward genetic approach using several types of pea mutants impaired in axillary bud outgrowth (Beveridge, 2000). Reciprocal grafting experiments with wild type and mutants of pea and *Arabidopsis* revealed that this hormone is mainly produced in roots and transported to shoots (Beveridge, 2006; Dun *et al.*, 2009; Beveridge and Kyozuka, 2010). The most probable route of hormone transportation appeared to be the xylem, and indeed, a canonical SL orobanchol was detected in the xylem sap of tomato and *Arabidopsis* (Kohlen *et al.*, 2011; Kohlen *et al.*, 2012). However, no other laboratories could confirm this finding. We also collected relatively large volumes of xylem sap from several plant species but could not detect any known SLs in them. Furthermore, SLs fed to roots of rice plants were detected in the shoots harvested 20 h after treatment, but not in the xylem sap (Xie *et al.*, 2015). These results indicate that SLs are transported from roots to shoots relatively slowly, although not through the xylem, but probably through hypodermis passage cells as in petunias where polar and asymmetric localizations of an ABC transporter, *Petunia axillaris* PLEITROPIC DRUG RESISTANCE 1 (PaPDR1), have been shown to mediate directional SL transport as well as localized exudation into the rhizosphere (Kretzschmar *et al.*, 2012; Sasse *et al.*, 2015).

As mentioned, rice produces only orobanchol-type SLs but tobacco produces both orobanchol- and strigol-types. When roots of these plants were treated with a mixture of four stereoisomers of 5DS (5DS, 4DO, and their enantiomers), only the orobanchol-type SL, 4DO, was detected in shoots of rice plants. However, both orobanchol- and strigol-type SLs, 4DO and 5DS, were detected in shoots of tobacco plants harvested 20 h after treatment, indicating that root to shoot transport of SLs is a structure- and stereo-specific process (Xie *et al.*, 2016). Although root-applied strigol was not transported to shoots in rice plants, it did strongly inhibit rice tillering (Umehara *et al.*, 2008), suggesting that the shoot branching inhibiting hormone is different from known canonical SLs. As mentioned earlier, the limited distribution of canonical SLs in the plant kingdom (**Table 1**) also supports this hypothesis.

What then, is this true hormone molecule? It should be noted that CL and its oxidized metabolites are widely distributed in the plant kingdom. Therefore, it is likely that these non-canonical SLs or their further metabolites function as plant hormones regulating shoot branching and other biological processes. For example, although root-applied strigol is not transported to shoots in rice plants, it may elicit release of hormone(s) which move shootward from cell to cell or through the xylem. This hypothesis can explain why root-fed GR5 (**32**), a synthetic SL containing only the C-D ring moiety, was highly active in inhibition of shoot branching but only weakly active when applied to the axillary buds of *Arabidopsis* (Umehara *et al.*, 2015). This implies that root-fed GR5 itself was not transported to the buds but GR5 strongly elicited release of the true hormone or signaling molecule(s) which moved upward to shoots. Unfortunately, in addition to the apparent scarcity, chemical instability makes isolation and structural determination of true hormone molecule(s) difficult. Furthermore, exogenous application of these molecules may not rescue a SL-deficient phenotype due to their instabilities. It is likely however that feeding experiments of candidate compounds or their precursors may provide insight into the identity of the hormone inhibiting the shoot branching.

## SLs as germination stimulants for root parasitic plants

It is well known that SLs were originally identified as germination stimulants for root parasitic plants: witchweeds (*Striga* spp.) and broomrapes (*Orobanche* and *Phelipanche* spp.). Canonical and non-canonical SLs elicit *Striga* and *Orobanche* seed germination at as low as pM to µM concentrations. The seeds of various *Striga* and *Orobanche* species show different sensitivities to each SL (Kim *et al.*, 2010; Kisugi *et al.*, 2013) and plants in general do not release a single SL but a mixture of at least two SLs. For example, 11 different canonical SLs were detected in root exudate of tobacco (Xie *et al.*, 2013; Xie, 2016). In addition to these canonical SLs, tobacco exudes the non-canonical SL, CLA. Therefore, it is likely that not a single SL but a SL mixture profile in the root exudate contributes to host-specific germination of root parasitic plant seeds. So far, effects of SL mixtures on seed germination stimulation have not been reported. There may be additive, synergistic, and antagonistic effects in mixtures of SLs on their germination stimulation activities. For example, root exudates of legumes that are good SL producers did not induce seed germination of *O. cumana*, indicating that an antagonistic effect occurs among SLs produced by leguminous plants on *O. cumana* seed germination (Fernández-Aparicio *et al.*, 2011). This suggests that seed germination of a particular root parasite may be reduced or inhibited through modifications of the SL profile of a crop as in *Striga*-resistant sorghum cultivars (Gobena *et al.*, 2017). In contrast, fractions from reverse phase high-performance LC separation of root exudates from English ivy (*Hederae helix*) that were active for *O. hederae* germination were not active for *O. minor* germination and vice versa (Kim *et al.*, unpublished), indicating the presence of host-specific germination stimulants.

## SLs as host recognition signals for AM fungi

Structural requirements for hyphal branching activity of canonical (Akiyama *et al.*, 2010) and some non-canonical SLs (Mori *et al.*, 2016) have been reported. In general, canonical SLs are more active than non-canonical SLs in inducing hyphal branching in *Gigaspora margarita*. Although CL is a weakly active branching factor, oxidation at C_19_ greatly enhanced its potency and CLA was as active as the typical canonical SL strigol (Mori *et al.*, 2016). Since CLA has been detected in root exudates from various plant species, CLA can be considered the most common branching factor released from plant roots (Yoneyama *et al.*, 2017). Then the question arises – why did plants develop canonical SLs? The chemical instability of CLA could be one reason, as it may decompose too rapidly to be sensed by AM fungi. In addition, CLA as a free acid may move and leach in soil more easily than do canonical SLs. These would have hampered precise host recognition by AM fungi and thus resulted in the evolution of canonical SLs. Characterization of SL receptor(s) in AM fungi and also in root nodule bacteria will shed light on why plants produce various canonical and non-canonical SLs.

## Conclusion

The wider distribution of non-canonical SLs, in particular CLA, in plant root exudates than that of canonical SLs suggests that CLA and some non-canonical SLs may be more common than canonical SLs as rhizosphere signals. Although we reported detection of orobanchol in *Arabidopsis* root exudates (Goldwasser *et al.*, 2008), we could not confirm this with our new MS system. Germination stimulants released from *Arabidopsis* roots (Goldwasser *et al.*, 2000) may be CLA and its derivatives rather than canonical SLs. This in turn implies that soil microorganisms which have been exposed to these signaling molecules for 400 million years would utilize them as an energy source and modify them metabolically, in some cases, into more stable and active compounds like canonical SLs. Germination stimulation by rhizosphere soils would be due in part to these metabolites (Zhang *et al.*, 2013). In addition, both canonical and non-canonical SLs and other short-lived rhizosphere signals would also be involved in plant–plant communications including self/non-self and kin-recognitions (Falik *et al.*, 2003; Biedrzycki *et al.*, 2014).

## Acknowledgements

This work was supported by the Science and Technology Research Promotion Program for Agriculture, Forestry, Fisheries and Food Industry; by KAKENHI (15K07093 and 16H04914); and by a grant from the JGC-S Scholarship Foundation to XX. Kaori Yoneyama is supported by the RPD project (JSPS).

## References

Abe S, Sado A, Tanaka K, Kisugi T, Asami K, Ota S, Kim HI, Yoneyama K, Xie X, Ohnishi T, Seto Y, Yamaguchi S, Akiyama K, Yoneyama K, Nomura T. 2014. Carlactone is converted to carlactonoic acid by MAX1 in *Arabidopsis* and its methyl ester can directly interact with AtD14 in vitro. Proceedings of the National Academy of Sciences of the United States of America 111, 18084–18089.

Akiyama K, Matsuzaki K, Hayashi H. 2005. Plant sesquiterpenes induce hyphal branching in arbuscular mycorrhizal fungi. Nature 435, 824–827.

Akiyama K, Ogasawara S, Ito S, Hayashi H. 2010. Structural requirements of strigolactones for hyphal branching in AM fungi. Plant and Cell Physiology 51, 1104–1117.

Al-Babili S, Bouwmeester HJ. 2015. Strigolactones, a novel carotenoid-derived plant hormone. Annual Review of Plant Biology 66, 161–186.

Alder A, Jamil M, Marzorati M, Bruno M, Vermathen M, Bigler P, Ghisla S, Bouwmeester H, Beyer P, Al-Babili S. 2012. The path from β-carotene to carlactone, a strigolactone-like plant hormone. Science 335, 1348–1351.

Bertin C, Yang X, Weston LA. 2003. The role of root exudates and allelochemicals in the rhizosphere. Plant and Soil 67, 67–83.

Beveridge CA. 2000. Long-distance signalling and a mutational analysis of branching in pea. Plant Growth Regulation 32, 193–203.

Beveridge CA. 2006. Axillary bud outgrowth: sending a message. Current Opinion in Plant Biology 9, 35–40.

Beveridge CA, Kyozuka J. 2010. New genes in the strigolactone-related shoot branching pathway. Current Opinion in Plant Biology 13, 34–39.

Biedrzycki ML, Jilany TA, Dudley SA, Bais HP. 2014. Root exudates mediate kin recognition in plants. Communicative & Integrative Biology 3, 28–35.

Boyer F-D, de Saint Germain A, Pillot J-P, Pouvreau J-B, Chen VX, Ramos S, Stévenin A, Simier P, Delavault P, Beau J-M, Rameau C. 2012. Structure-activity relationship studies of strigolactone-related molecules for branching inhibition in garden pea: molecule design for shoot branching. Plant Physiology 159, 1524–1544.

Boyer F-D, de Saint Germain A, Pouvreau JB, Clave G, Pillot JP, Roux A, Rasmussen A, Depuydt S, Lauressergues D, Frei Dit Frey N, Heugebaert TS, Stevens CV, Geelen D, Goormachtig S, Rameau C. 2014. New strigolactone analogs as plant hormones with low activities in the rhizosphere. Molecular Plant 7, 675–690.

Charnikhova TV, Gaus K, Lumbroso A, Sanders M, Vincken J-P, De Mesmaeker A, Ruyter-Spira CP, Screpanti C, Bouwmeester HJ. 2017. Zealactones. Novel natural strigolactones from maize. Phytochemistry 137, 123–131.

Chen VX, Boyer F-D, Rameau C, Retailleau P, Vors J-P, Beau J-M. 2010. Stereochemistry, total synthesis, and biological evaluation of the new plant hormone solanacol. Chemistry 16, 13941–13945.

Cohen M, Prandi C, Occhiato EG, Tabasso S, Wininger S, Resnick N, Steinberger Y, Koltai H, Kapulnik Y. 2013. Structure–function relations of strigolactone analogs: activity as plant hormones and plant interactions. Molecular Plant 6, 141–152.

Cook CE, Whichard LP, Turner B, Wall ME, Egley GH. 1969. Germination of witchweed (*Striga lutea* Lour.): isolation and properties of a potent stimulant. Science 154, 1189–1190.

Dun EA, Brewer PB, Beveridge CA. 2009. Strigolactones: discovery of the elusive shoot branching hormone. Trends in Plant Science 14, 364–372.

Falik O, Reides P, Gersani M, Novoplansky A. 2003. Self/non-self discrimination in roots. Journal of Ecology 91, 525–531.

Fernández-Aparicio M, Yoneyama K, Rubiales D. 2011. The role of strigolactones in host specificity of *Orobanche* and *Phelipanche* seed germination. Seed Science Research 21, 55–61.

Flematti GR, Scaffidi A, Waters MT, Smith SM. 2016. Stereospecificity in strigolactone biosynthesis and perception. Planta 243, 1361–1373.

Fukui K, Ito S, Asami T. 2013. Selective mimics of strigolactone actions and their potential use for controlling damage caused by root parasitic weeds. Molecular Plant 6, 88–99.

Gobena D, Shimels M, Rich PJ, Ruyter-Spira C, Bouwmeester H, Kanuganti S, Mengiste T, Ejeta G. 2017. Mutation in sorghum *LOW GERMINATION STIMULANT 1* alters strigolactones and causes *Striga* resistance. Proceedings of the National Academy of Sciences of the United States of America 114, 4471–4476.

Goldwasser Y, Plakhine D, Yoder JI. 2000. *Arabidopsis thaliana* susceptibility to *Orobanche* spp. Weed Science 48, 342–346.

Goldwasser Y, Yoneyama K, Xie X, Yoneyama K. 2008. Production of strigolactones by *Arabidopsis thaliana* responsible for *Orobanche aegyptiaca* seed germination. Plant Growth Regulation 55, 21–28.

Gomez-Roldan V, Fermas S, Brewer PB, Puech-Pagès V, Dun EA, Pillot J-P, Letisse F, Matusova R, Danoun S, Portais J-C, Bouwmeester H, Bécard G, Beveridge CA, Rameau C, Rochange SF. 2008. Strigolactone inhibition of shoot branching. Nature 455, 189–194.

Hauck C, Müller S, Schildknecht H. 1992. A germination stimulant for parasitic flowering plants from *Sorghum bicolor*, a genuine host plant. Journal of Plant Physiology 139, 474–478.

Jamil M, Charnikhova T, Houshyani B, van Ast A, Bouwmeester HJ. 2011. Genetic variation in strigolactone production and tillering in rice and its effect on *Striga hermonthica* infection. Planta 235, 473–484.

Jamil M, Kanampiu FK, Karaya H, Charnikhova T, Bouwmeester HJ. 2012. *Striga hermonthica* parasitism in maize in response to N and P fertilisers. Field Crops Research 134, 1–10.

Kim HI, Kisugi T, Khetkam P, Xie X, Yoneyama K, Uchida K, Yokota T, Nomura T, McErlean CS, Yoneyama K. 2014. Avenaol, a germination stimulant for root parasitic plants from *Avena strigosa*. Phytochemistry 103, 85–88.

Kim HI, Xie X, Kim HS, Chun JC, Yoneyama K, Nomura T, Takeuchi Y, Yoneyama K. 2010. Structure-activity relationship of naturally occurring strigolactones in *Orobanche minor* seed germination stimulation. Journal of Pesticide Science 35, 344–347.

Kisugi T, Xie X, Kim HI, Yoneyama K, Sado A, Akiyama K, Hayashi H, Uchida K, Yokota T, Nomura T, Yoneyama K. 2013. Strigone, isolation and identification as a natural strigolactone from *Houttuynia cordata*. Phytochemistry 87, 60–64.

Kohlen W, Charnikhova T, Lammers M, Pollina T, Toth P, Haider I, Pozo MJ, de Maagd RA, Ruyter-Spira C, Bouwmeester HJ, Lopez-Raez JA. 2012. The tomato *CAROTENOID CLEAVAGE DIOXYGENASE8 (SlCCD8)* regulates rhizosphere signaling, plant architecture and affects reproductive development through strigolactone biosynthesis. New Phytologist 196, 535–547.

Kohlen W, Charnikhova T, Liu Q, Bours R, Domagalska MA, Beguerie S, Verstappen F, Leyser O, Bouwmeester H, Ruyter-Spira C. 2011. Strigolactones are transported through the xylem and play a key role in shoot architectural response to phosphate deficiency in nonarbuscular mycorrhizal host Arabidopsis. Plant Physiology 155, 974–987.

Kondo Y, Tadokoro E, Matsuura M, Iwasaki K, Sugimoto Y, Miyake H, Takikawa H, Sasaki M. 2007. Synthesis and seed germination stimulating activity of some imino analogs of strigolactones. Bioscience, Biotechnology, and Biochemistry 71, 2781–2786.

Kretzschmar T, Kohlen W, Sasse J, Borghi L, Schlegel M, Bachelier JB, Reinhardt D, Bours R, Bouwmeester HJ, Martinoia E. 2012. A petunia ABC protein controls strigolactone-dependent symbiotic signalling and branching. Nature 483, 341–344.

Mori N, Nishiuma K, Sugiyama T, Hayashi H, Akiyama K. 2016. Carlactone-type strigolactones and their synthetic analogues as inducers of hyphal branching in arbuscular mycorrhizal fungi. Phytochemistry 130, 90–98.

Nefkens GHL, Thuring JWJF, Beenakkers MFM, Zwanenburg B. 1997. Synthesis of a phthaloylglycine-derived strigol analogue and its germination stimulatory activity toward seeds of the parasitic weeds *Striga hermonthica* and *Orobanche crenata*. Journal of Agricultural and Food Chemistry 45, 2273–2277.

Proust H, Hoffmann B, Xie X, Yoneyama K, Schaefer DG, Yoneyama K, Nogué F, Rameau C. 2011. Strigolactones regulate protonema branching and act as a quorum sensing-like signal in the moss *Physcomitrella patens*. Development 138, 1531–1539.

Sasse J, Simon S, Gübeli C, Liu G-W, Cheng X, Friml J, Bouwmeester H, Martinoia E, Borghi L. 2015. Asymmetric localizations of the ABC transporter PaPDR1 trace paths of directional strigolactone transport. Current Biology 25, 647–655.

Sato D, Awad AA, Takeuchi Y, Yoneyama K. 2005. Confirmation and quantification of strigolactones, germination stimulants for root parasitic plants *Striga* and *Orobanche*, produced by cotton. Bioscience Biotechnology and Biochemistry 69, 98–102.

Screpanti C, Fonné-Pfister R, Lumbroso A, Rendine S, Lachia M, De Mesmaeker A. 2016. Strigolactone derivatives for potential crop enhancement applications. Bioorganic & Medicinal Chemistry Letters 26, 2392–2400.

Seto Y, Sado A, Asami K, Hanada A, Umehara M, Akiyama K, Yamaguchi S. 2014. Carlactone is an endogenous biosynthetic precursor for strigolactones. Proceedings of the National Academy of Sciences of the United States of America 111, 1640–1645.

Siame BA, Weerasuriya Y, Wood K, Ejeta G, Butler LG. 1993. Isolation of strigol, a germination stimulant for *Striga asiatica,* from host plants. Journal of Agricultural and Food Chemistry 41, 1486–1491.

Thuring JWJF, Nefkens GHL, Zwanenburg B. 1997. Asymmetric synthesis of all stereoisomers of the strigol analogue GR24. Dependence of absolute configuration on stimulatory activity of *Striga hermonthica* and *Orobanche crenata* seed germination. Journal of Agricultural and Food Chemistry 45, 2278–2283.

Tokunaga T, Hayashi H, Akiyama K. 2015. Medicaol, a strigolactone identified as a putative didehydro-orobanchol isomer, from *Medicago truncatula*. Phytochemistry 111, 91–97.

Ueno K, Furumoto T, Umeda S, Mizutani M, Takikawa H, Batchvarova R, Sugimoto Y. 2014. Heliolactone, a non-sesquiterpene lactone germination stimulant for root parasitic weeds from sunflower. Phytochemistry 108, 122–128.

Umehara M, Cao M, Akiyama K, Akatsu T, Seto Y, Hanada A, Li W, Takeda-Kamiya N, Morimoto Y, Yamaguchi S. 2015. Structural requirements of strigolactones for shoot branching inhibition in rice and Arabidopsis. Plant and Cell Physiology 56, 1059–1072.

Umehara M, Hanada A, Yoshida S, Akiyama K, Arite T, Takeda-Kamiya N, Magome H, Kamiya Y, Shirasu K, Yoneyama K, Kyozuka J, Yamaguchi S. 2008. Inhibition of shoot branching by new terpenoid plant hormones. Nature 455, 195–200.

Xie X. 2016. Structural diversity of strigolactones and their distribution in the plant kingdom. Journal of Pesticide Science 41, 175–180.

Xie X, Kisugi T, Yoneyama K, Nomura T, Akiyama K, Uchida K, Yokota T, McErlean CSP, Yoneyama K. 2017. Methyl zealactonoate, a novel germination stimulant for root parasitic weeds produced by maize. Journal of Pesticide Science 42, 58–61.

Xie X, Kusumoto D, Takeuchi Y, Yoneyama K, Yamada Y, Yoneyama K. 2007. 2’-Epi-orobanchol and solanacol, two unique strigolactones, germination stimulants for root parasitic weeds, produced by tobacco. Journal of Agricultural and Food Chemistry 55, 8067–8072.

Xie X, Yoneyama K, Harada Y, Fusegi N, Yamada Y, Ito S, Yokota T, Takeuchi Y, Yoneyama K. 2009a. Fabacyl acetate, a germination stimulant for root parasitic plants from *Pisum sativum*. Phytochemistry 70, 211–215.

Xie X, Yoneyama K, Kisugi T, Nomura T, Akiyama K, Asami T, Yoneyama K. 2015. Strigolactones are transported from roots to shoots, although not through the xylem. Journal of Pesticide Science 40, 214–216.

Xie X, Yoneyama K, Kisugi T, Nomura T, Akiyama K, Asami T, Yoneyama K. 2016. Structure- and stereospecific transport of strigolactones from roots to shoots. Journal of Pesticide Science 41, 55–58.

Xie X, Yoneyama K, Kisugi T, Uchida K, Ito S, Akiyama K, Hayashi H, Yokota T, Nomura T, Yoneyama K. 2013. Confirming stereochemical structures of strigolactones produced by rice and tobacco. Molecular Plant 6, 153–163.

Xie X, Yoneyama K, Kurita J-y, Harada Y, Yamada Y, Takeuchi Y, Yoneyama K. 2009b. 7-Oxoorobanchyl acetate and 7-oxoorobanchol as germination stimulants for root parasitic plants from flax (*Linum usitatissimum*). Bioscience, Biotechnology and Biochemistry 73, 1367–1370.

Xie X, Yoneyama K, Kusumoto D, Yamada Y, Takeuchi Y, Sugimoto Y, Yoneyama K. 2008. Sorgomol, germination stimulant for root parasitic plants, produced by Sorghum bicolor. Tetrahedron Letters 49, 2066–2068.

Xie X, Yoneyama K, Yoneyama K. 2010. The strigolactone story. Annual Review of Phytopathology 48, 93–117.

Yasui M, Ota R, Tsukano C, Takemoto Y. 2017. Total synthesis of avenaol. Nature Communications 8, 674. doi: 10.1038/s41467-017-00792-1.

Yokota T, Sakai H, Okuno K, Yoneyama K, Takeuchi Y. 1998. Alectrol and orobanchol, germination stimulants for *Orobanche minor*, from its host red clover. Phytochemistry 49, 1967–1973.

Yoneyama K, Arakawa R, Ishimoto K, Kim HI, Kisugi T, Xie X, Nomura T, Kanampiu F, Yokota T, Ezawa T, Yoneyama K. 2015. Difference in *Striga*-susceptibility is reflected in strigolactone secretion profile, but not in compatibility and host preference in arbuscular mycorrhizal symbiosis in two maize cultivars. New Phytologist 206, 983–989.

Yoneyama K, Awad AA, Xie X, Yoneyama K, Takeuchi Y. 2010. Strigolactones as germination stimulants for root parasitic plants. Plant and Cell Physiology 51, 1095–1103.

Yoneyama K, Mori N, Sato T, XIe X, Ohnishi T, Yokota T, Akiyama K, Yoneyama K, Nomura T. 2017. Conversion of carlactone to carlactonoic acid is a common and conserved function of MAX1 homologs in plants. New Phytologist, submitted.

Yoneyama K, Xie X, Kusumoto D, Sekimoto H, Sugimoto Y, Takeuchi Y, Yoneyama K. 2007. Nitrogen deficiency as well as phosphorus deficiency in sorghum promotes the production and exudation of 5-deoxystrigol, the host recognition signal for arbuscular mycorrhizal fungi and root parasites. Planta 227, 125–132.

Yoneyama K, Xie X, Sekimoto H, Takeuchi Y, Ogasawara S, Akiyama K, Hayashi H, Yoneyama K. 2008. Strigolactones, host recognition signals for root parasitic plants and arbuscular mycorrhizal fungi, from Fabaceae plants. New Phytologist 179, 484–494.

Yoneyama K, Xie X, Yoneyama K, Takeuchi Y. 2009. Strigolactones: structures and biological activities. Pest Management Science 65, 467–470.

Zhang W, Ma Y, Wang Z, Ye X, Shui J. 2013. Some soybean cultivars have ability to induce germination of sunflower broomrape. PLos One 8, e59715.

Zhang Y, van Dijk ADJ, Scaffidi A, Flematti GR, Hofmann M, Charnikhova T, Verstappen F, Hpeworth J, van der Krol S, Leyser O, Smith SM, Zwanenburg B, Al-Babili S, Ruyter-Spira C, Bouwmeester HJ. 2014. Rice cytochrome P450 MAX1 homologs catalyze distinct steps in strigolactone biosynthesis. Nature Chemical Biology 10, 1028–1033.

Zwanenburg B, Mwakaboko AS, Reizelman A, Anilkumar G, Sethumadhavan D. 2009. Structure and function of natural and synthetic signalling molecules in parasitic weed germination. Pest Management Science 65, 478–491.

